# Approximating Scaffold Printability Utilizing Computational Methods

**DOI:** 10.1101/2022.07.11.499589

**Authors:** Ashkan Sedigh, Pejman Ghelich, Jacob Quint, Mohamadmahdi Samandari, Ali Tamayol, Ryan E. Tomlinson

**Affiliations:** Department of Orthopaedic Surgery, Thomas Jefferson University, Philadelphia, PA; Department of Biomedical Engineering, University of Connecticut, Farmington, CT

**Keywords:** Extrusion Bioprinting, Fuzzy System, Adaptive Neuro Fuzzy System, Gelatin, Approximation

## Abstract

Bioprinting facilitates the generation of complex, three-dimensional (3D), cell-based constructs for various applications. Although multiple bioprinting technologies have been developed, extrusion-based systems have become the dominant technology due to the diversity of materials (bioinks) that can be utilized, either individually or in combination. However, each bioink has unique material properties and extrusion characteristics that affect bioprinting utility, accuracy, and precision. Here, we have extended our previous work to achieve high precision (i.e., repeatability) across samples by optimizing bioink-specific printing parameters. Specifically, we hypothesized that an adaptive neuro-fuzzy inference system (ANFIS) could be used as a computational method to address the imprecision in 3D bioprinting test data and uncover the optimal printing parameters for a specific bioink that result in high accuracy and precision. To test this hypothesis, we have implemented an ANFIS model consisting of four inputs (bioink concentration, printing pressure, speed, and temperature) and a single output to quantify the precision (scaffold bioprinted linewidth range). We validate our use of the bioprinting precision index (BPI) with both standard and normalized printability factors. In total, our results indicate that computational methods are a cost-efficient measure to improve the precision and robustness of extrusion 3D bioprinting with gelatin-based bioinks.

## 1. Introduction

### 1.1. Bioprinting

Additive manufacturing is a process by which three-dimensional objects are generated by material deposition in sequential layers [1]–[3]. Bioprinting is an emerging field of additive manufacturing, in which bioactive scaffolds can be quickly generated by the deposition of layers of cell-laden biocompatible materials, such as collagen or other hydrogels [4]. After slicing a solid model, the appropriate extrusion path and required printing parameters can be used to direct the fabrication of the construct in a layer-by-layer fashion. Indeed, the ability to place cells in biologically relevant scaffold materials with high spatial resolution has made bioprinting a popular fabrication method for tissue engineering [4]–[6].

However, the parameters used to perform the bioprinting procedure itself will have significant effects on the final properties of the model [7]. Therefore, it is essential to fully characterize and optimize the bioprinting parameters (e.g., print speed or bioink viscosity) necessary to reach the desired outputs, such as architectural matching of the defect shape, high cell viability, appropriate cell function, and mechanical properties [8]. For example, increasing the nozzle size on the bioprinter decreases the shear stress placed on the biomaterial during extrusion, resulting in increased cell viability but reduced print resolution [9]. Therefore, determining the optimal print parameters is imperative for success in bioprinting. As a result, several studies have been performed in the field of bioprinting optimization, such as optimization of a solid model for 3D bioprinting, bioink optimization, and bioprinting parameter selection [10], [11].

Hydrogels provide three-dimensional (3D) support for cellular growth and tissue formation similar to native extracellular matrix (ECM) and are widely utilized to study cellular proliferation, migration, differentiation, and interaction [12]–[14]. In general, hydrogels are crosslinked networks of hydrophilic polymers that swell in water [13], [15], [16]. One of the most common natural hydrogels in biomedical applications and bioprinting is gelatin, which is derived through collagen hydrolysis. Although there are several ways to hydrolyze the collagen, we have used Type A (acid-derived) gelatin in this experiment. Gelatin is particularly useful for drug release and tissue engineering due to its biocompatibility [17], rapid biodegradability, constituent purity, high cell attachments due to arginine-glycine-aspartic acid (RGD) sequences [18], and tunable physical properties [16]. The composition, crosslinking method, and level of crosslinking of gelatin-based materials affect its physical and biochemical properties. Crosslinking gelatin bioink can be done by photo crosslinking [7], [19], enzymatic crosslinking [20], thermal crosslinking [21], or chemical crosslinking [22]. In this study, we used gelatin as our bioink with thermal crosslinking. As a result, our study is directly applicable to a variety of other gelatin-based bioinks, such as gelatin methacryloyl (GelMA) [13], [23].

### 1.2. Approximation and Prediction

There is a significant degree of imprecision and uncertainty inherent in bioprinting optimization. A potential approach to handling this issue, which arises from normal biological variation, is implementing approximation systems based on computational methods [24]. Systems biology has become a critical multidisciplinary research area between computer science and biology in recent years. Studies in this field aim to develop computational models of biological processes, requiring both a robust dynamic model and a large dataset of experimental results [25], [26].

Developing a dynamic model is challenging and requires well-characterized control parameters to approximate laboratory experiments’ outcomes. Nonetheless, recent studies have observed that machine learning algorithms can effectively predict output parameters using either a deterministic or stochastic biological model. A partial list of the approaches employed to this end includes meta-heuristic, evolutionary, global optimization, genetic programming, simulated annealing, simplex, ant-colony, fuzzy genetic hybrid system, and multi-objective optimization [25], [27]. In-silico models have helped reproducibility and quality in the in-vitro experiments significantly in tissue [28]-[30].

### 1.3 Adaptive Neuro-Fuzzy Inference System (ANFIS)

Here, we have developed a quantitative model of a biomanufacturing process using the ANFIS approach, which is a potential solution for overcoming uncertainty in an experimental dataset. Previous studies have shown that the accuracy of a fuzzy system approach is the same as the deterministic mathematical approach (ordinary differential equations) for the same kinetic dataset [31]. Moreover, fuzzy systems can be utilized to find the qualitative system response when a quantitative dataset is unavailable [32], [33]. In decision-making systems, theoretical fuzzy models have been used to approximate biomechanical properties such as stress/strain of a bone structure in a biological process [34]–[36]. The mathematical model of the ANFIS system is described in the supplementary section of the paper.

### 1.4 Approximation in 3D printed Scaffold

Bioprinting outputs can be predicted using classic physics formulation or computational and machine learning methods. We recently demonstrated that computational methods are precise in the output approximation, which can be measured and optimized based on the bioprinting inputs of speed, pressure, and dilution percentage [34]. Herein, we propose to implement an approximation model using a 3D scaffold with greater complexity of inputs. To do so, we implemented an ANFIS algorithm and measured the training error on the dataset. Ultimately, we optimized and approximated the bioprinting output for extrusion parameters with three different precision indexes and measured the testing error. Fig. 1 shows the general workflow of the process.

**Figure 1.**
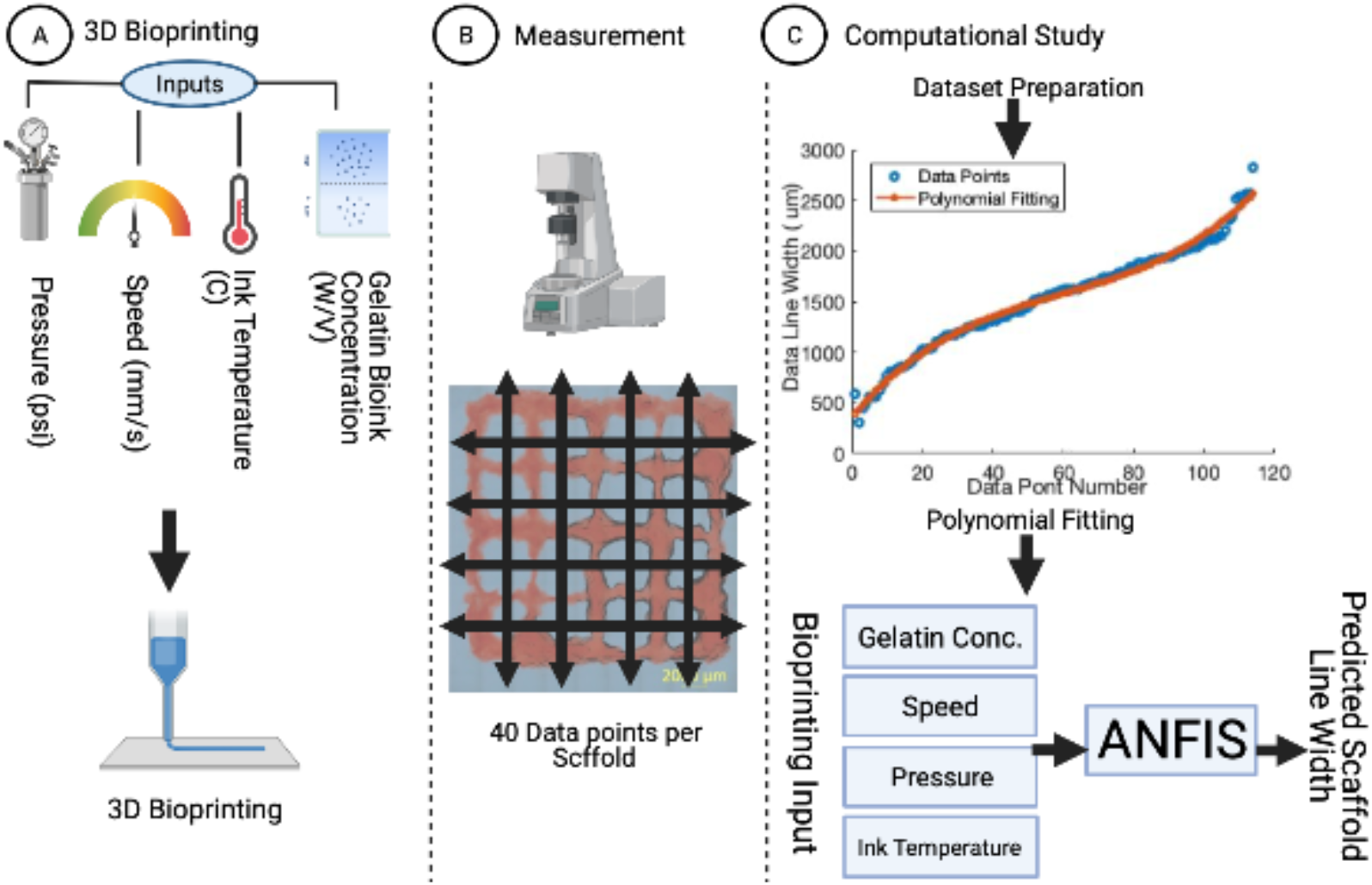
General workflow of the study. (A) First, the 3D model was designed with four different input parameters and the sample was printed by an extrusion-based bioprinter. (B) Next, the data was acquired using brightfield microscopy. (C) The data were normalized, processed, and interpolated by a polynomial function. Then the polynomial coefficients were used as the output for the training model; the final model was generated to calculate the BPI for the required bioprinting parameter’s set.

## 2. Methods

### 2.1. Bioprinting

Type A gelatin from porcine skin (Gel strength 300, Sigma, US, lot # SLCD2367) was measured and gradually added to warmed Dulbecco’s phosphate-buffered saline (DPBS) (no calcium, no magnesium, 14190136, Gibco) to give a 10% w/v stock solution. Also, 100 μL of red food dye was added to the stock for better contrast during the imaging. From the stock solution, three different concentrations were made (3%, 4%, and 5% w/v) and stored in 15 mL centrifuge tubes.

Next, 3 mL of each gelatin bioink was pipetted into a 5 mL syringe. A 22-gauge tapered plastic tip was used for bioprinting after being loaded into an Allevi® 3 3D bioprinter. The print layout was a 20 mm x 20 mm meshed square with a 500 μm layer height. Infill distances were 5 mm. The printability was investigated under various ranges of ink temperature (4 to 16 °C), ink concentration (3 to 5 % w/v), pressure (6 to 11 psi), and tip speed (2 to 10 mm/s) (Table 1), to provide a training data set for the predicting algorithm. Three replicates of each condition were performed (n=3). The bioinks were maintained at the printing temperature for at least 10 minutes before printing. The room humidity was maintained at less than 40% with a room temperature range of 22-24 °C. A stitched brightfield image of printed patterns from 5X magnified tiles was saved and postprocessed using a ZEISS Axio Observer microscope. Only data points described in Table 2 were successfully 3D printed in the experimental step.

**Table 1.** Experiment Design

| Run | Gelatin<br>Bioink Conc.<br>(w/v %)<br>MF SD: 4.24 | Printing Speed (mm/s)<br>MF SD: 0.42 | Pressure (PSI) | Bioink Temp. (C)<br>MF SD:16.99 |
| --- | --- | --- | --- | --- |
| 1 | 3 | 6 | 8.5 | 10 |
| 2 | 4 | 6 | 6 | 10 |
| 3 | 4 | 6 | 8.5 | 10 |
| 4 | 3 | 10 | 6 | 16 |
| 5 | 3 | 10 | 11 | 16 |
| 6 | 3 | 2 | 11 | 4 |
| 7 | 5 | 10 | 6 | 16 |
| 8 | 3 | 2 | 6 | 16 |
| 9 | 5 | 2 | 11 | 16 |
| 10 | 5 | 2 | 6 | 4 |
| 8.5 | 4 | 6 | 8.5 | 10 |
| 10 | 4 | 6 | 8.5 | 16 |
| 13 | 5 | 6 | 8.5 | 10 |
| 14 | 3 | 10 | 11 | 4 |
| 15 | 3 | 2 | 6 | 4 |
| 11 | 4 | 6 | 8.5 | 10 |
| 17 | 4 | 6 | 8.5 | 10 |
| 18 | 5 | 2 | 6 | 16 |
| 19 | 5 | 10 | 11 | 4 |
| 16 | 4 | 6 | 8.5 | 10 |
| 21 | 5 | 2 | 11 | 4 |
| 22 | 5 | 10 | 11 | 16 |
| 23 | 4 | 6 | 8.5 | 10 |
| 24 | 5 | 10 | 6 | 4 |
| 25 | 4 | 6 | 8.5 | 10 |
| 26 | 4 | 6 | 8.5 | 4 |
| 27 | 4 | 6 | 11 | 10 |
| 28 | 3 | 10 | 6 | 4 |
| 29 | 4 | 10 | 8.5 | 10 |
| 30 | 3 | 2 | 11 | 16 |
| 31 | 4 | 2 | 8.5 | 10 |

**Table 2.**
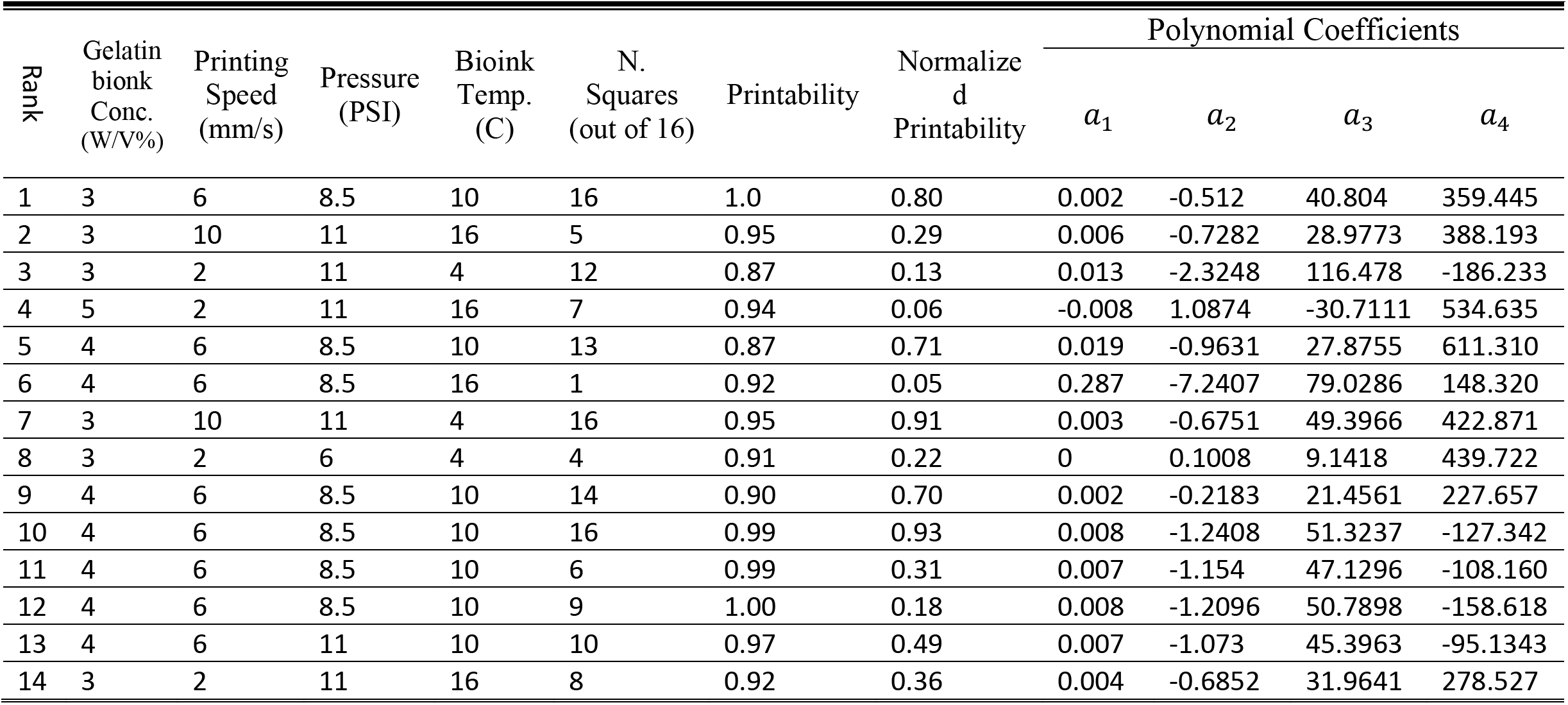
ANFIS Training dataset and Printability Factor

### 2.2. Imaging and Analysis

Each printed sample was imaged using brightfield microscopy 5X magnified tiles and analyzed in FIJI [21]. To determine and quantify the actual bioprinted construct line width, we measured each side of the squares (16 squares in each sample – 96-point data). Each parameter set was bioprinted in three individual runs. All three runs were pooled into one array and utilized in the fitting function interpolation.

### 2.3. Polynomial Approximation

In order to train the ANFIS with our data series for each bioprinted scaffold, we used polynomial fitting to find the representation for each data series. The measured data can be compacted and expanded with a proper fitting function. Since the data points can be expanded with the known polynomial coefficients, we utilize the fitted polynomial coefficients as the inputs for the ANFIS training. To find the optimal output data representatives in the learning process, we compare the Root Mean Squared Error (RMSE) for four standard fitting functions (Figure 3). We used the below fitting function to evaluate our printed data series:

3-degree polynomial (four variables):

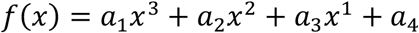

5-degree polynomial (six variables):

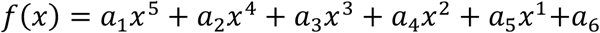

2-term Exponential (four variables):

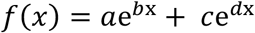

2-term Gaussian (six variables):

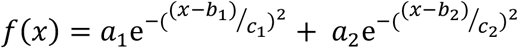

Fig. 3 shows the fitting rate of the tested interpolation functions (2-term Gaussian, 3-Degree Polynomial, 5-Degree Polynomial, 2-term exponential). The RMSE value between interpolation functions was not significant, so we utilized the function with the simplest and the fewest number of coefficients (third-degree polynomial). The fitting evaluation was then performed on the experimental data from the 30 bioprinted scaffolds.

### 2.3. Implementing Adaptive Neuro Fuzzy Inference System

The ANFIS structure is programmed utilizing the MATLAB ANFIS editor toolbox to train the ANFIS model and approximate the polynomial coefficients. The Sugeno-based fuzzy inference system and the Gaussian membership functions are used to train the datasets and map the relationship between process inputs and the polynomial coefficients (see Supplemental Methods). In the first step, we trained ANFIS with four inputs (gelatin concentration (% w/v), bioink temperature (°C), pressure (psi), tip speed (mm/s)), and output (polynomial coefficients). Table 2 indicates the training dataset, including the four inputs and four outputs (coefficients). In the next step, we measured the error between the expected line width and the predicted polynomials. Ultimately, we used the introduced adaptive neuro-fuzzy system to optimize the precision in three different grades. We considered errors between 0-400 μm to be high precision, 500-1000 μm to be medium precision, and 1000-2500 μm to be low precision [34], [36]. Fig. 2 shows the implemented ANFIS with inputs/outputs and the ANFIS layers.

**Figure 2.**
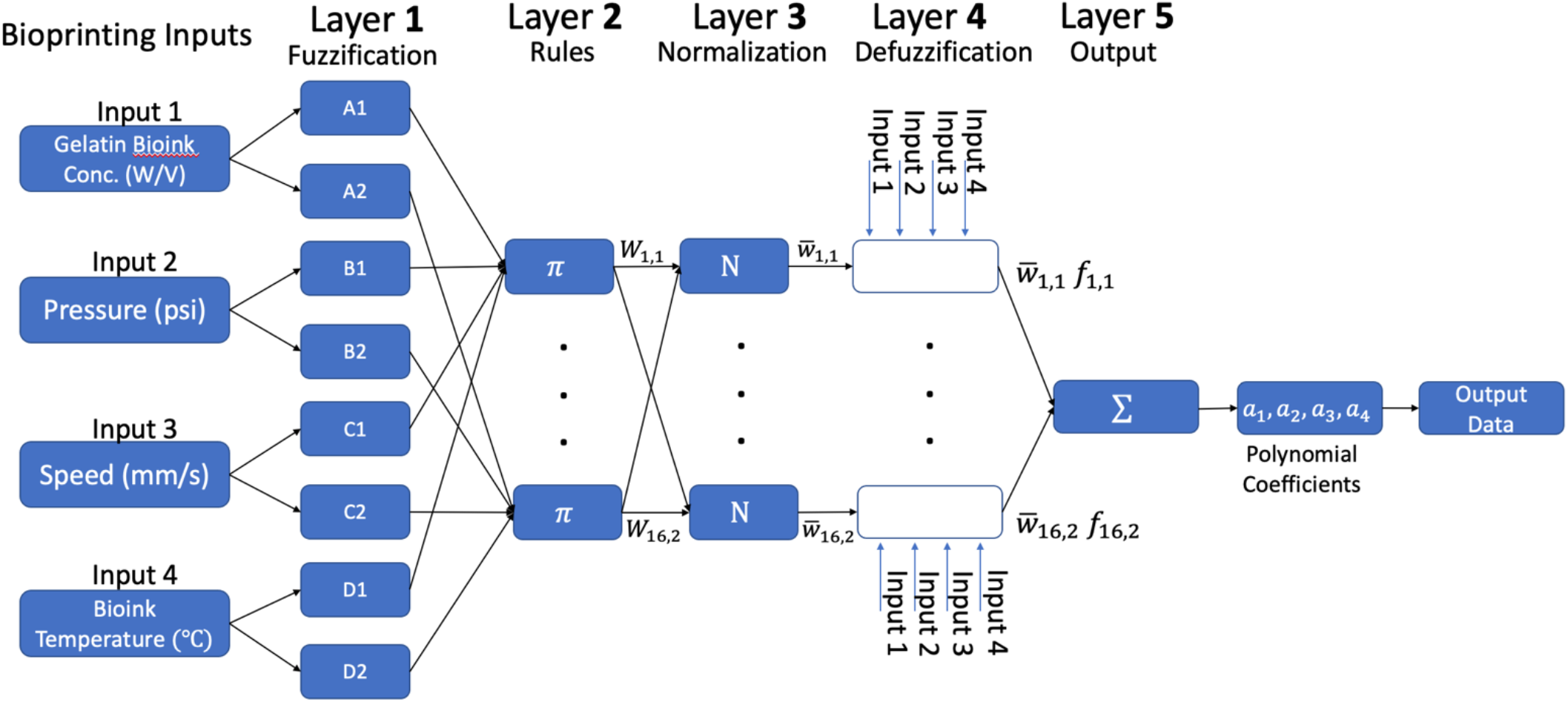
ANFIS structure and study design. A general overview and different processing layers of the ANFIS including fuzzification, rules, normalization, defuzzification, and output. It includes four inputs (gelatin bioink concentration, speed, pressure, and bioink temperature) and polynomial coefficients.In the next step, the line width is approximated with the predicted polynomial coefficients.

**Figure 3.**
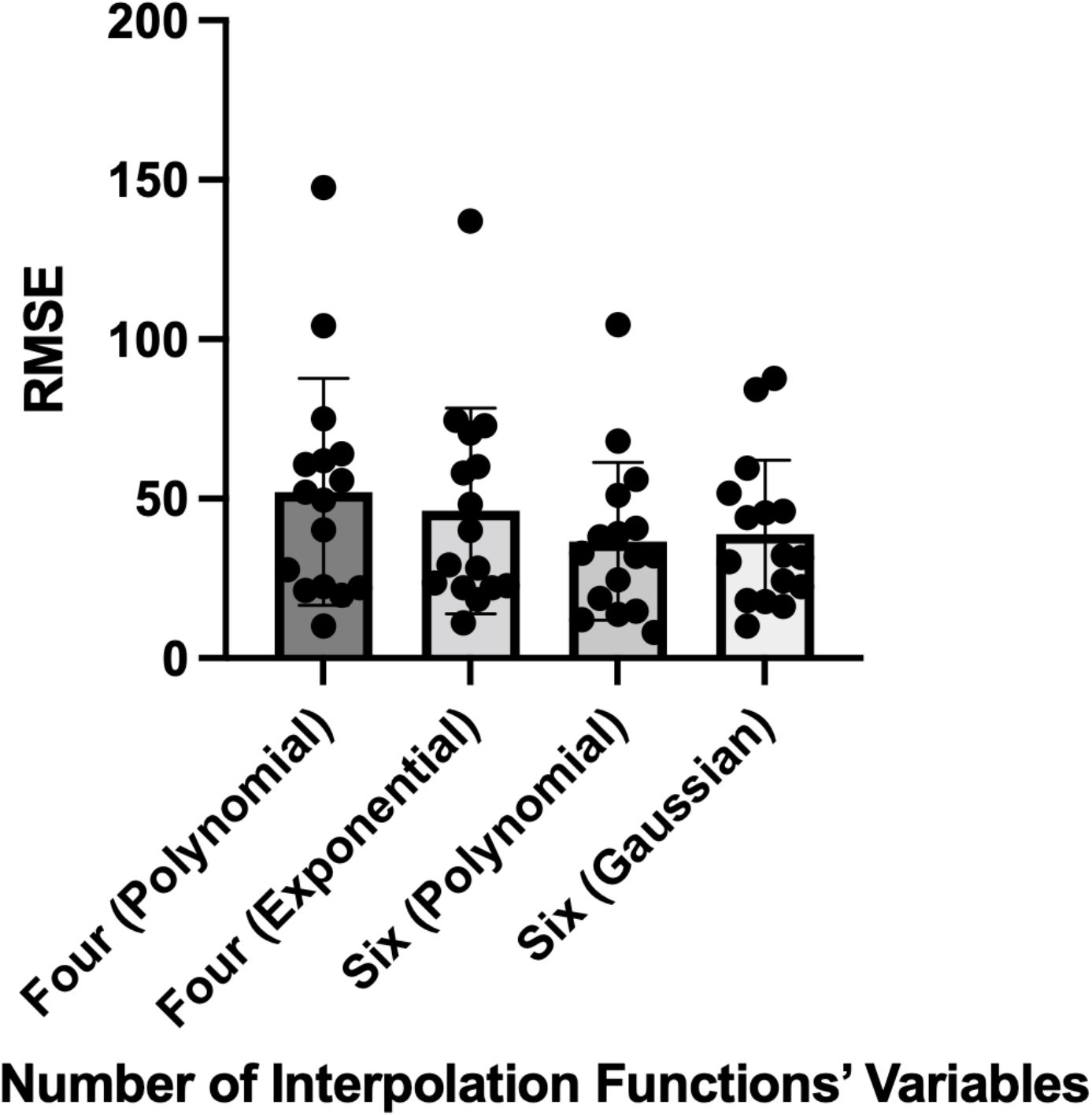
Evaluation of multiple interpolation functions. In this experiment, we examined multiple interpolation functions as the ANFIS output model. There is no significant difference between discussed interpolation functions RMSE.

### 2.4. Bioprinting Precision Index

In recent work utilizing a fuzzy system approach to bioprinting parameter optimization [34], we introduced the Bioink Precision Index (BPI) as a new metric for evaluating bioink precision. BPI is defined as the gradient of the fuzzy 3D surfaces. The standard calculation for a gradient of a 3D surface with four inputs:

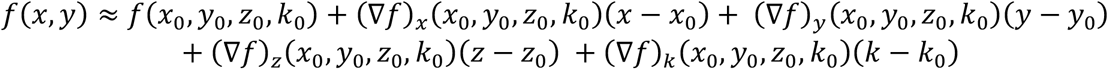

Thus, BPI is calculated from the sum of the squared error of the above equation:

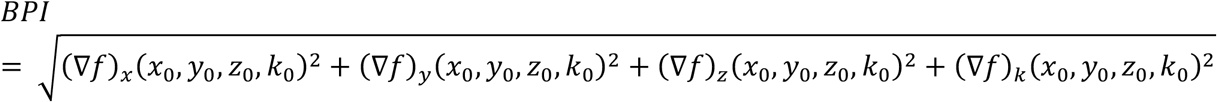

Where *x*_0_, *y*_0_, *z*_0_, and *k*_0_ are the inputs in our system (gelatin concentration, pressure, speed, and temperature).

### 2.5. Bioprintibility Factor

Bioprintability is a factor that quantifies shape fidelity with a referenced formula for multiple shapes. For a square shape, the bioink printability (*pr*) uses the following function:

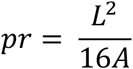

Where *L* is the perimeter and *A* is the area of the enclosed area. For an ideal gelation condition or perfect printability status, the interconnected channels of the constructs would demonstrate a square shape, and the *pr* value is 1. *pr* indicates the degree of gelatin bioink to maintain the construct shape fidelity [37]. We use the following normalization coefficient to normalize the *pr* value:

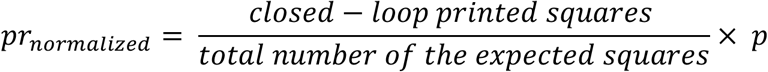

## 3. Results

### 3.1. Principal Component Analysis Indicates the Influential Printing Parameters

Our system has four inputs (gelatin bioink concentration, speed, pressure, and bioink temperature) with four outputs (*a*_4_, *a*_2_, *a*_3_, *a*_4_), so we used Principal Component Analysis (PCA) with Alternating Least Squares (ALS) to find the component vectors. Here, the first two components cover 80 percent of the data in our analysis (Fig. 4 shows the training and testing data on a PCA biplot). Principal Component 1 (PC1) and Principal Component 2 (PC2) explanatory forms are shown in the below equations:

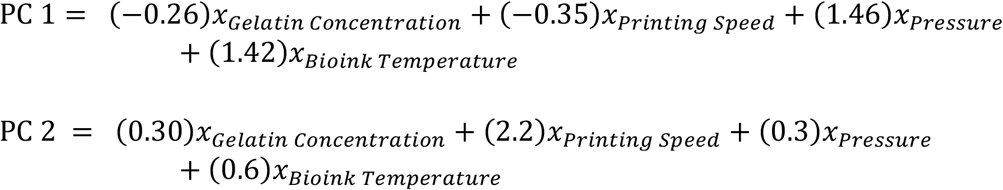

**Figure 4.**
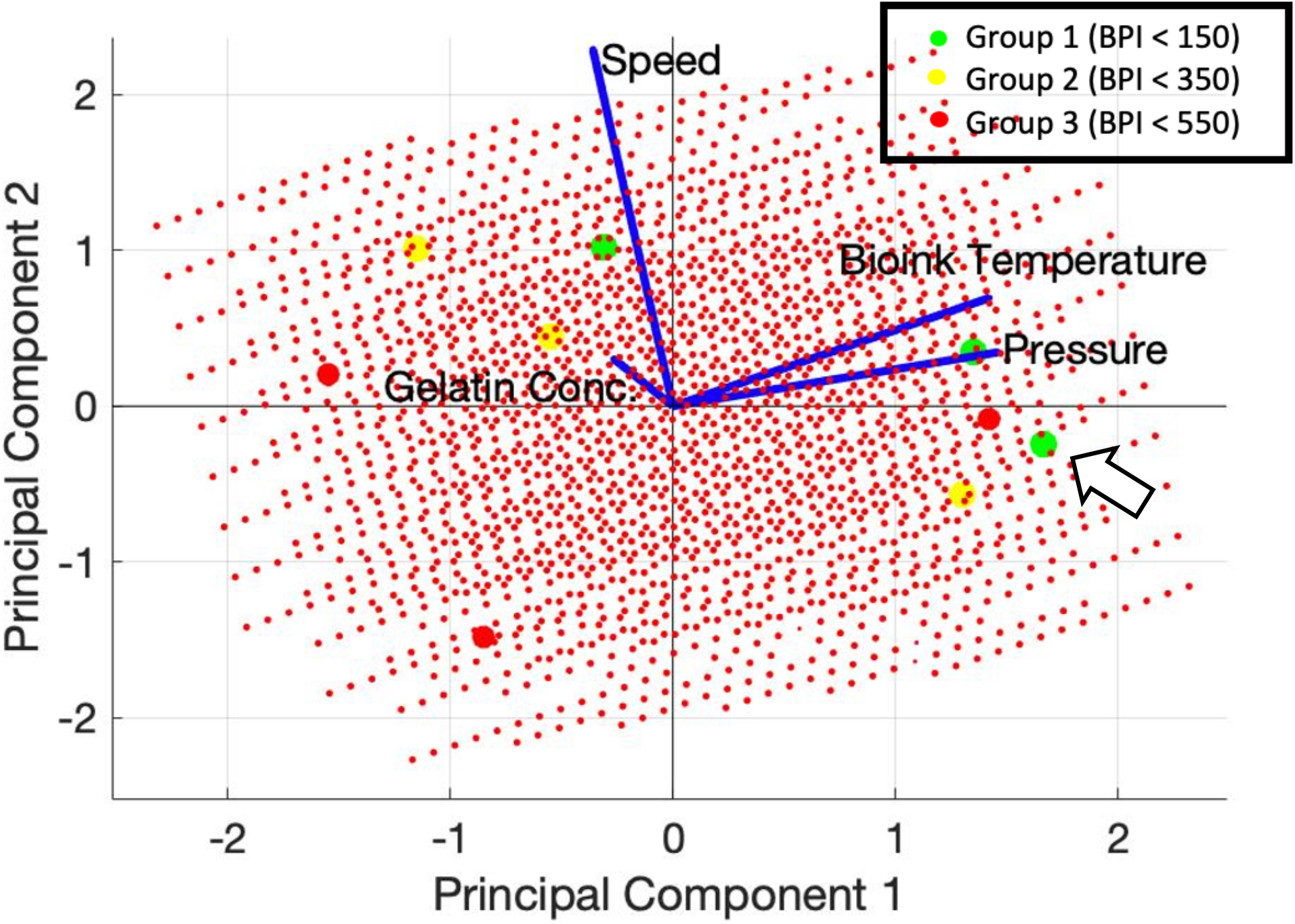
PCA biplot with training and testing data. We use PCA to lower the input dimension to be able to visualize it. In this biplot, the small red dots are the training inputs, and the (green, yellow, and red) points are the testing data. The approximated BPI for the testing points is shown.

PCA biplot indicates that the bioink temperature and pressure parameters are highly and positively correlated in the First PC. However, the speed parameter is highly and positively correlated to the second PC. Gelatin concentration has the lowest correlation in the first and second PCs. This result is consistent with the PC1 and PC2 explanatory equations indicating that the gelatin concentration has the lowest coefficients. Therefore, we conclude that gelatin concentration has the lowest and negative correlation and could be omitted in the experiment.

### 3.2. Training ANFIS

Next, we trained our ANFIS model with the prepared dataset (Table 2). The dataset includes gelatin concentration (% w/v), pressure (psi), tip speed (mm/s), and temperature (°C). In this model, we use the polynomial coefficients as the training output (*a*_1_, *a*_2_, *a*_3_, *a*_4_). ANFIS algorithm predicted the coefficients for the data that had not been in the training dataset. The training process with the provided dataset (Table 2) resulted in an RMSE value of 0.008 - *a*_1_ 0.55 - *a*_2_, 13.97 - *a*_3_, and 182.07 - *a*_4_ (Fig. 5). S. Fig. 3 shows the microscopic images of the extruded training dataset. Next, we approximated the fuzzy output for the data between the minimum and maximum of each parameter as determined by the physical limitations of the bioprinter. Fig. 6 shows the 3D plot of PC1 and PC2 for each coefficient as an output (*a*_1_, *a*_2_, *a*_3_, *a*_4_). Consequently, the final scaffold width measurements can be extracted by replacing the approximated four coefficients in the polynomial function within the predefined data range. In the final step, we calculate the BPI value for the data points with the given description to find the minimum BPI, which represents parameter sets with maximum precision and bioprintability.

**Figure 5.**
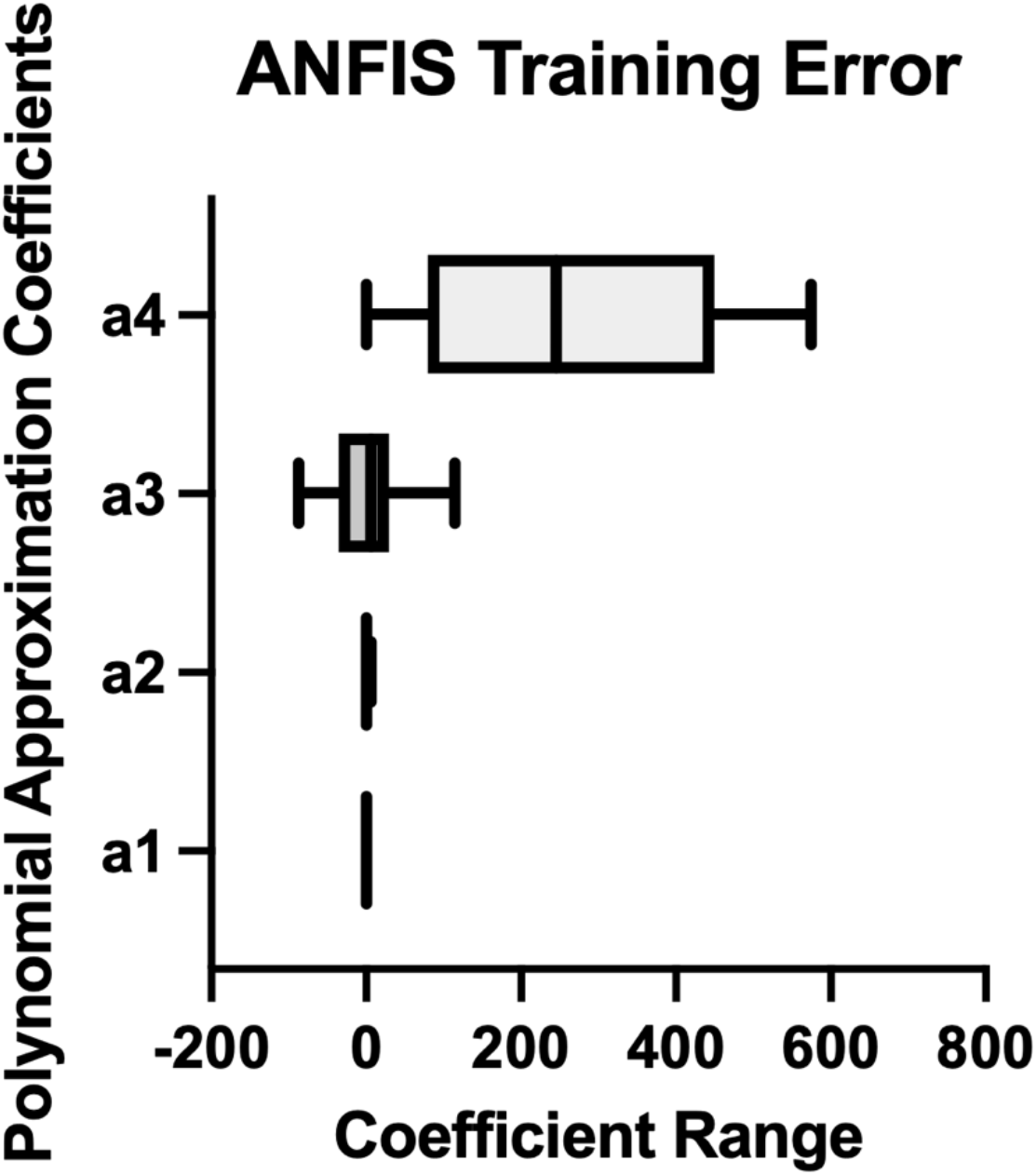
ANFIS training error. The RMSE error for the training dataset with four coefficient outputs is indicated. has the highest rank in the interpolation function, and it is trained with the highest accuracy, whereas *a*_4_ has the lowest rank and highest training error.

**Figure 6.**
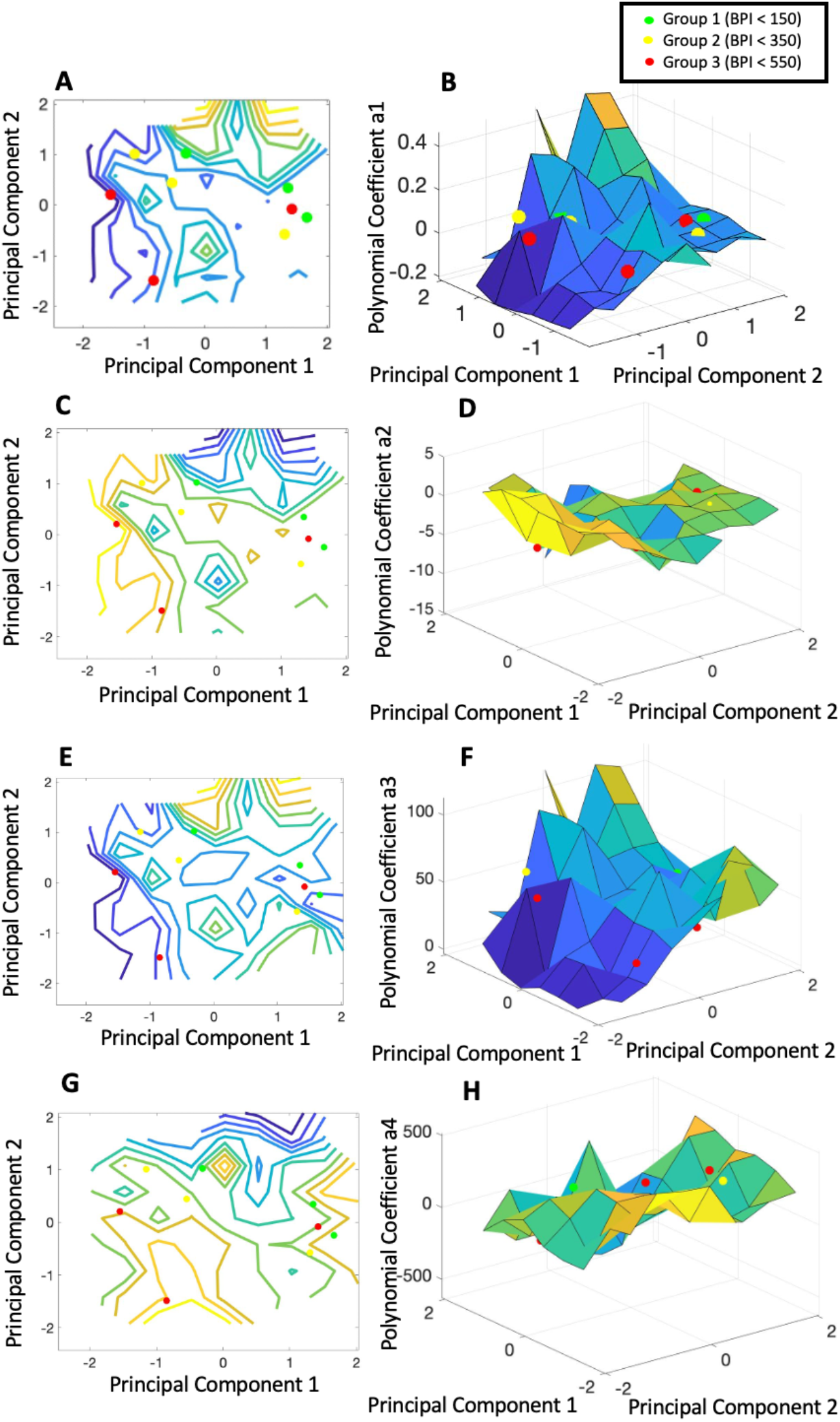
Fuzzy approximation of training and testing data. The general shape of the 3D graphs of Gelatin bioink shows that the multi-input multi-output system is a non-convex function with multiple optimal points. Here, the slightest slope results in a higher BPI. Coefficient *a*_3_ has the minimum effect on the output since it has a flat surface.

### 3.3. Testing ANFIS and BPI Approximation

To test our implemented model for optimizing BPI and validating that the minimum BPI values in our algorithm are applicable at the practical level, we categorized BPI into three levels (0-150, 150-350, and greater than 350). We randomly chose the inputs that fit the three groups that were not in the training set. As a result, Fig. 4 shows the testing values in three groups on the PC1-PC2 biplot (0-150 Green, 150-350 Yellow, and 350-above Red). Particularly, the green data points show the minimum BPI and highest precision and robustness, whereas the red data points show the lowest precision. Fig. 6 shows the testing data points on the approximated coefficient 3D plots. We printed the testing data (Table 3) in the same conditions as the training set (Fig. 7). Each parameter set was repeated in three individual runs. As a result, one of the bioprinting parameter sets did not meet the expectation (Fig 7-c).

**Table 3.** Testing Groups Experiment Design

| Run | Gelatin Bionk<br>Concentration<br>(W/V%) | Printing<br>Speed<br>(mm/s) | Pressure (PSI) | Bioink<br>Temperature (C) |
| --- | --- | --- | --- | --- |
| Group 1<br>BPI: 0 - 150 |  |  |  |  |
| 1 | 4 | 6 | 11 | 10 |
| 2 | 4 | 9 | 8 | 10 |
| 3 | 4 | 4 | 11 | 12 |
| Group 2<br>BPI: 150 - 350 |  |  |  |  |
| 4 | 3 | 5 | 10 | 11 |
| 5 | 5 | 6 | 8 | 10 |
| 6 | 5 | 8 | 7 | 10 |
| Group 3<br>BPI: 350 - above |  |  |  |  |
| 7 | 3 | 4 | 6 | 8 |
| 8 | 5 | 3 | 11 | 13 |
| 9 | 5 | 6 | 6 | 10 |

**Figure 7.**
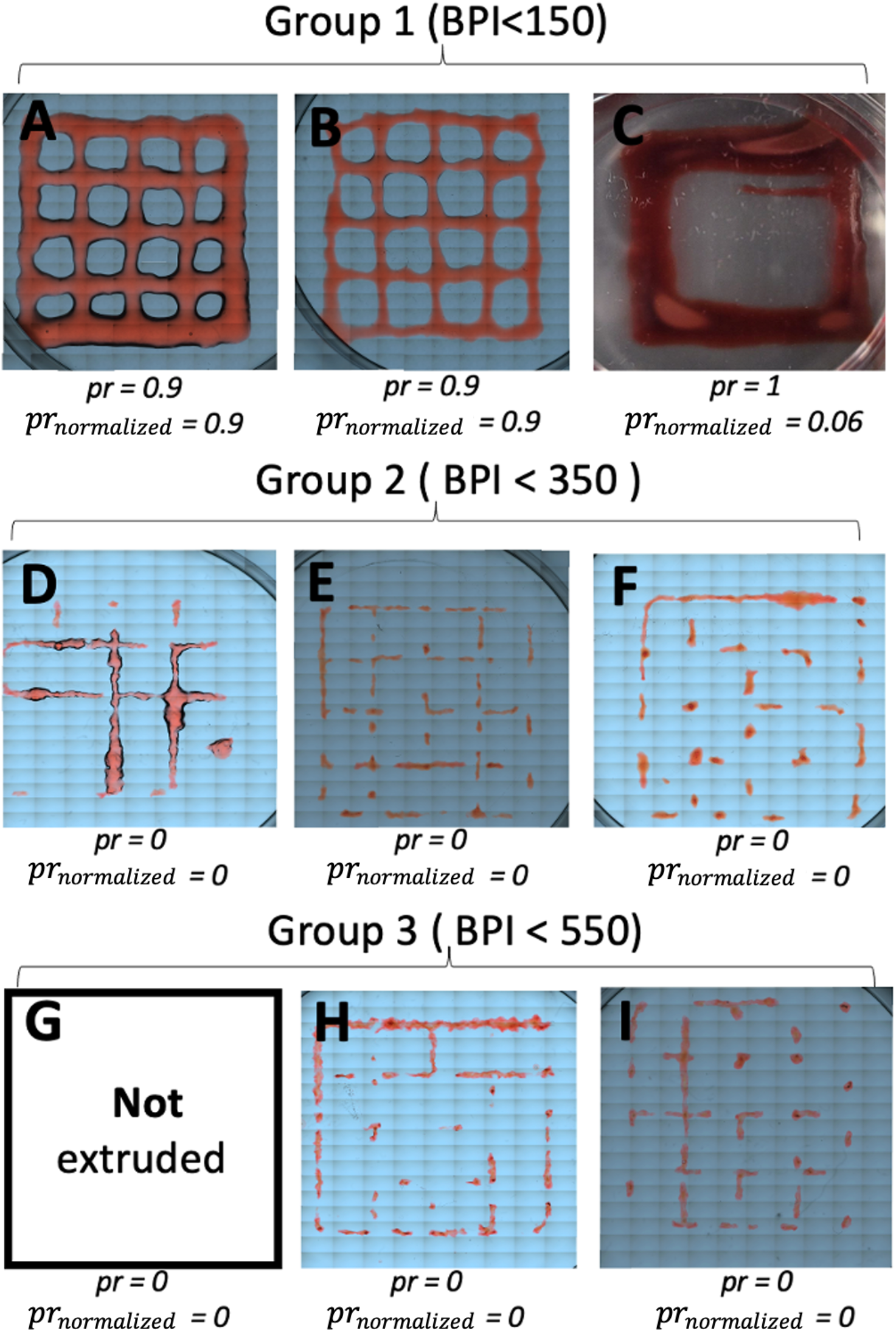
Bioprinted test data. The test data in three groups were 3D printed with the same environmental condition as the training group. The first group has the highest precision (Lowest BPI), the middle group has the medium and the last group has the lowest precision (Higher BPI). The printability factor (*pr*) and (*pr_normalized_*) for each printing parameter set is presented.

### 3.4 Comparison of Bioprintibility Factor and Normalized Bioprintibility Factor

Table 2 shows the calculated *pr* value for the training dataset. However, *pr* value does not include the generalization term to calculate the bioprintibility for the entire construct. For instance, in the experiment, constructs may have 8 squares with *pr* = 1, but the other half may not be printed or with a very low *pr* <0.5 value. Since the *pr* value does not show a significant resolution to interpret data, we used the weighted method to normalize the *pr* value for the bioprinted scaffold. In our experiment, each sample has 16 squares. The Normalized Bioprintability Factor result is shown in Table 2, which increased the resolution and the data significance in terms of interpretation and further comparison. T-test between Standard Bioprintability factor (*pr*) and the Normalized Bioprintability Factor (Table 2) revealed a statistically significant difference (p < 0.001).

## 4. Discussion

In this study, we introduced a novel approach to approximate the Bioink Precision Index (BPI) in 3D by utilizing an adaptive neuro-fuzzy interference system (ANFIS) in the multiple input-output bioprinting process. In the training model, the inputs include gelatin bioink concentration (W/V%), bioprinting pressure (psi), speed (mm/s), and bioink temperature (C) with one output - construct line width (mm), which is derived from predicted thepolynomial coefficients. Next, the BPI surface was generated using the training data. We then tested three independent bioprinting parameter sets that had not been in the training dataset from high, medium, and low BPI (9 total). Perhaps surprisingly, the input data for these parameter sets had a very narrow range for each input. In particular, gelatin bioink concentration ranged from 3-5%, speed from 2 to 10 mm/s, pressure from 6 to 11 psi, and temperature from 4 to 16 °C. Nonetheless, the output and bioprintability of these parameter sets varied widely – from flawless to useless. Moreover, the experimental results were consistent with the approximation (BPI) that accurately predicted the high, medium, and low precision. In total, our study demonstrates that computational methods are a cost-effective way to significantly improve precision in 3D bioprinting.

The difference between fuzzy systems and machine learning methods such as neural networks is that fuzzy system is a decision-making process from raw and ambiguous data, whereas neural networks rely on training data to learn and improve system accuracy over time. Therefore, by utilizing the hybrid model (ANFIS), we are able to train a fuzzy system to be used in inferencing systems where data vagueness is the primary obstacle. Moreover, ANFIS systems have transparency in data inferencing and approximation, whereas the inference model cannot be precisely extracted using machine learning methods. Here, we use the ANFIS model to increase the precision and maintain a robust approximator. We also note that the learning feature can be used to adapt our system for most bioinks and bioprinting environments. Importantly, conventional physics-based formulations cannot be utilized for optimization since they are not reversible in the multi-parameter domain. Also, the non-convex solution space and discrete values in the traditional formulation of the process decrease the probability of finding the optimal and precise bioprinting solutions. Therefore, the optimization may result in limited optimal solutions (data points).

In our testing experiment, one of the three constructs printed with a low BPI (high precision) parameter set did not meet the precision requirement. Although the data point (indicated with an arrow in Fig. 4) was predicted to have high precision, it was not printable in the application. According to the PCA biplot (Fig. 4), the faulty data point is close to the high BPI (low precision) data points, whereas the others are close to the PCA vectors. As a result, we hypothesize that the faulty data point resulted from a fitting error in the multi-dimensional space with a narrow input data range (Fig. 7C). The fitting performance could be improved by utilizing a larger training dataset within the same input data range. Nonetheless, optimization without prior knowledge in a narrow data range and multi-dimensional space requires a soft computing approach to find high-precision solutions.

Bioprintibility factor (*pr*) is a metric previously established to quantify the bioprinted construct shape fidelity [37]. Although a shape can have the *pr* value equivalent to 1, this metric does not represent the overall 3D construct shape fidelity. Therefore, we introduced a new *pr_normalized_* metric generated by multiplying by the normalization coefficient. Fig. 7c shows a construct with the *pr* value equal to 1, which is inconsistent with the poor bioprintability (only 1 out of 16 squares extruded). After normalization, the value for this sample was reduced to *pr_normalized_* = 0.06, consistent with the actual result. Therefore, we have calculated the *number of printed squares, pr_normalized_*, and *pr* for the training dataset, which is available in Table 2.

One of the limitations of this study is the lack of experiments on multiple nozzle gauges. Since the BPI is specifically a measure of output precision, utilizing a parameter set with optimal BPI may not result in high accuracy. This limitation may result in a necessary tradeoff between accuracy and precision in choosing a parameter set for bioprinting. Future work may extend this approach to other parameters that affect bioprinting or its experimental outcomes, such as cell viability or biocompatibility. Nonetheless, we note that BPI is independent of input dimensions/units and will remain a valuable metric for comparison between bioinks as bioprinting technology evolves.

## 5. Conclusion

Obtaining high precision in bioprinting is a necessary step toward the mass production of bioprinted 3D constructs for use in research and medicine. Here, we have demonstrated that the Adaptive Neuro-Fuzzy Inference System (ANFIS) approach can be used to approximate output precision with a set of bioprinting parameters, including printing speed, extrusion pressure, gelatin bioink concentration, and bioink temperature, as well as determine bioprinting parameter sets that maximize precision in 3D constructs. Furthermore, we have applied the previously defined Bioink Precision Index (BPI) that can be used to quickly compare the ease of reproducibility regardless of the number of inputs, and environment.

## Acknowledgment

This research is supported by the National Institute of Arthritis and Musculoskeletal and Skin Diseases and the National Institute of Dental and Craniofacial Research of the National Institutes of Health under award numbers AR074953 (RET) and DE028397 (RET). Other financial support from the National Institutes of Health (AR077132, AR073822, AR079114) is gratefully acknowledged. The content is solely the responsibility of the authors and does not necessarily represent the official views of the funding bodies.

## Supplementary

Statistical Inferencing is a method to use a sample from a population to conclude about the entire population. Inferencing can be performed with three standard methods: Estimating, Prediction, and tolerance interval. The validity of inference is related to the way the data are obtained and the stationarity of the process producing the data. For instance, the least square method is one of the estimation models. Tolerance intervals are used to give a range of values that, with pre-specified confidence, will contain at least a pre-specified proportion of the measurements in the population.

One consideration in designing an experiment or sampling study is the desired precision utilizing estimators or predictors. The precision of an estimator is a measure of the estimator variability. Moreover, another equivalent way of expressing precision is the width of a level L confidence interval. For a given population, precision is a function of the sample size: the larger the sample, the greater the precision. However, if our population is not fixed, the precision of the output system is related to the system robustness. One of the methods in approximating the output with high precision, is the Fuzzy Systems.

The fuzzy system inherently doesn’t have the learning feature, The neural network provides the capability to adapt and learn to the fuzzy system. The Neural Network is utilized to tune the Fuzzy membership functions. Tuning membership functions helps to adjust the system more accurately based on the training dataset and reduces the cost of computational m Since the fuzzy system doesn’t have the learning feature, The neural network provides the capability to adapt and learn to the fuzzy system. The Neural Network is utilized to tune the Fuzzy membership functions. Tuning membership functions helps to adjust the system more accurately based on the training dataset and reduces the cost of computational model implementation. ANFIS consists of five layers to train, fuzzify, and defuzzify the output into ordinary numbers. In the supplementary section, we described the formulation and implementation of the ANFIS algorithm. ANFIS structure depends on essential parameters such as number of membership functions, number of iterations, type of membership function, etc.odel implementation. ANFIS consists of five layers to train, fuzzify, and defuzzify the output into ordinary numbers. In the supplementary section, we described the formulation and implementation of the ANFIS algorithm. ANFIS structure depends on essential parameters such as number of membership functions, number of iterations, type of membership function, etc.

Fuzzy logic is an extended model of standard logic. In standard logic, truth values can only be either completely false or completely true (with degrees of truth equal to 0 or 1, respectively), whereas, in fuzzy logic, values can have a degree of truth between 0 and 1. This generalization provides a mathematical framework to move from discrete to continuous values. In other words, in contrast to sets in classical logic, a fuzzy set is a set without a crisp boundary. For instance, if the reference set *X* is a Universe of discourse for elements *x*, the fuzzy set A is defined:

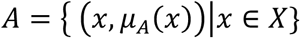

Where *μ_A_* (*x*) is called the Membership Function (MF) for the fuzzy set A. The MF maps each element of the Universe set *X* to a grade between 0 and 1, i.e., a membership of 0 means that the associated element is not included, whereas a membership 1 means a fully included element.

A fuzzy rule-based system is a modeling framework that uses the above fuzzy set theory along with a set of “if-then” rules where the antecedents and consequents are fuzzy logic propositions. This rule-based fuzzy system is used for modelling the inputs and their relationships with the output variables. A type-1 Fuzzy System (T1 FS) is a framework consisting of weighted rules, membership functions, and a fuzzy inference system. This system takes the crisp data (fuzzy singletons) or fuzzy inputs and generates fuzzy outputs based on the given if-then rules. A method of defuzzification is then used to extract a crisp value inferred from the fuzzy model.

From the commonly used membership functions such as triangular, trapezoidal and Gaussian, this study employs Gaussian membership function in ANFIS model for assessment of input and output variables. Further steps deal with providing the Gaussian membership functions along with fuzzy input to the neural network.

The back propagation algorithm is applied to train the inference engine for appropriate rule base selection in the neural network block. After adequate training, suitable rules are fired from the neural network to produce the optimal output (Table 2). Finally, the output variable in linguistic terms is converted into crisp output by the de-fuzzifier block. The structure of ANFIS based on Takagi–Sugeno system comprises of five different network layers, i.e., Layer 1 to Layer 5. For describing the ANFIS procedure, two inputs *x* and *y*, one output *f*. The Takagi-Sugeno ANFIS rules base consists of IF–THEN fuzzy rules and that can be expressed as:

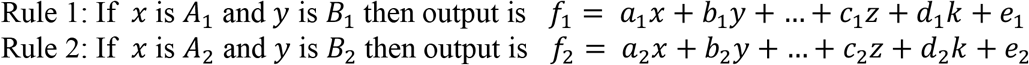

## Layer 1

The first layer deals with conversion of set of inputs (*x, y*) into linguistic terms (*A*_1_, *A*_2_, *B*_2_, *B*_1_ and *B*_2_) by applying membership function. Fuzzy layer comprises of set of adaptive nodes whose function is expressed as:

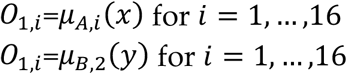

Where *O*_1,*i*_ represents the output function. *μ_A,i_*(*x*) and *μ_B,i_*(*y*) represents the membership functions. For the present study, Gaussian membership function is selected for input and triangular membership function is chosen for output.

## Layer 2

The second layer shown with circle deals with obtaining the output signal by multiplying the input signal received from previous layers with fixed node function denoted as *O*_2,*i*_. The output of every node provides the firing strength of a rule.

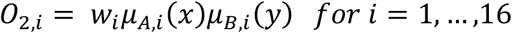

## Layer 3

The third layer also named as normalized layer, is represented as ratio of individual rule of firing strength to the algebraic sum of all the firing strength.

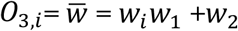

where *O*_3,*i*_ and 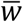 represents the output of third layer and firing strength, respectively.

## Layer 4

The fourth layer also referred to as defuzzification layer where each node of layer is adjustable and variable in nature. The output node function can be represented as:

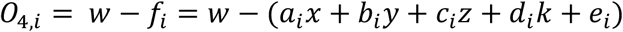

where *a_i_*, *b_i_*, *c_i_, d_i_* and *e_i_* are the set of consequent parameters for rule *l_i_*

## Layer 5

The fifth layer shown by circle consist of only one node. It evaluates the overall ANFIS output by summing up the all the output coming from previous layer and can be expressed as:

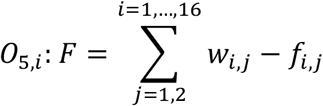

In the next step, we plot each approximated polynomial coefficient utilizing ANFIS with the PC1 and PC2 value (S. Fig. 1-2). The training samples microscopic images are shown S. Fig. 3. The bioprining properties for each sample is described in Table 2.

**Supplemental Figure 1.**
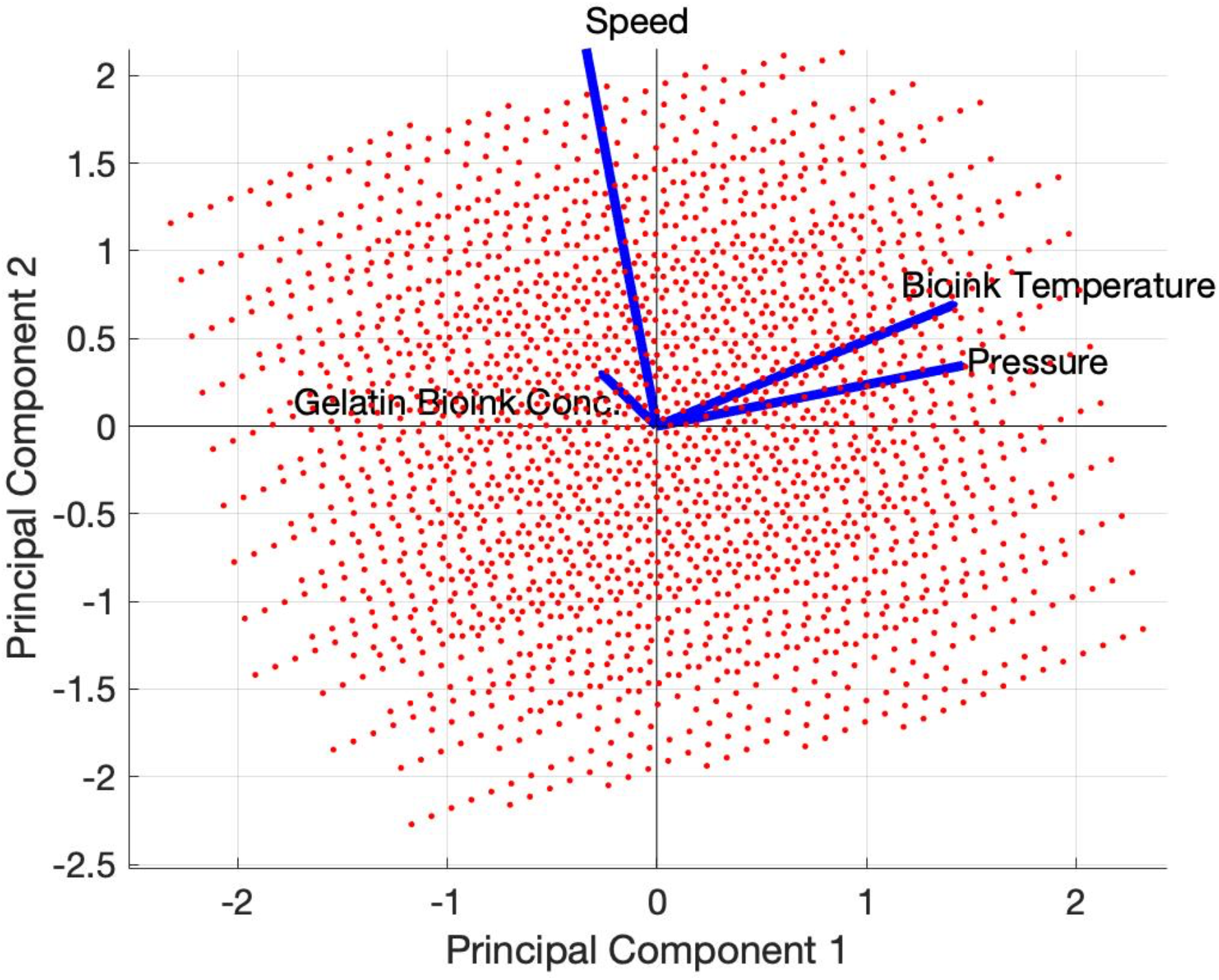
Principal Component 1 and 2 biplot. The biplot shows the correlation between the four input parameters in the first two components (80 percent data points coverage).

**Supplemental Figure 2.**
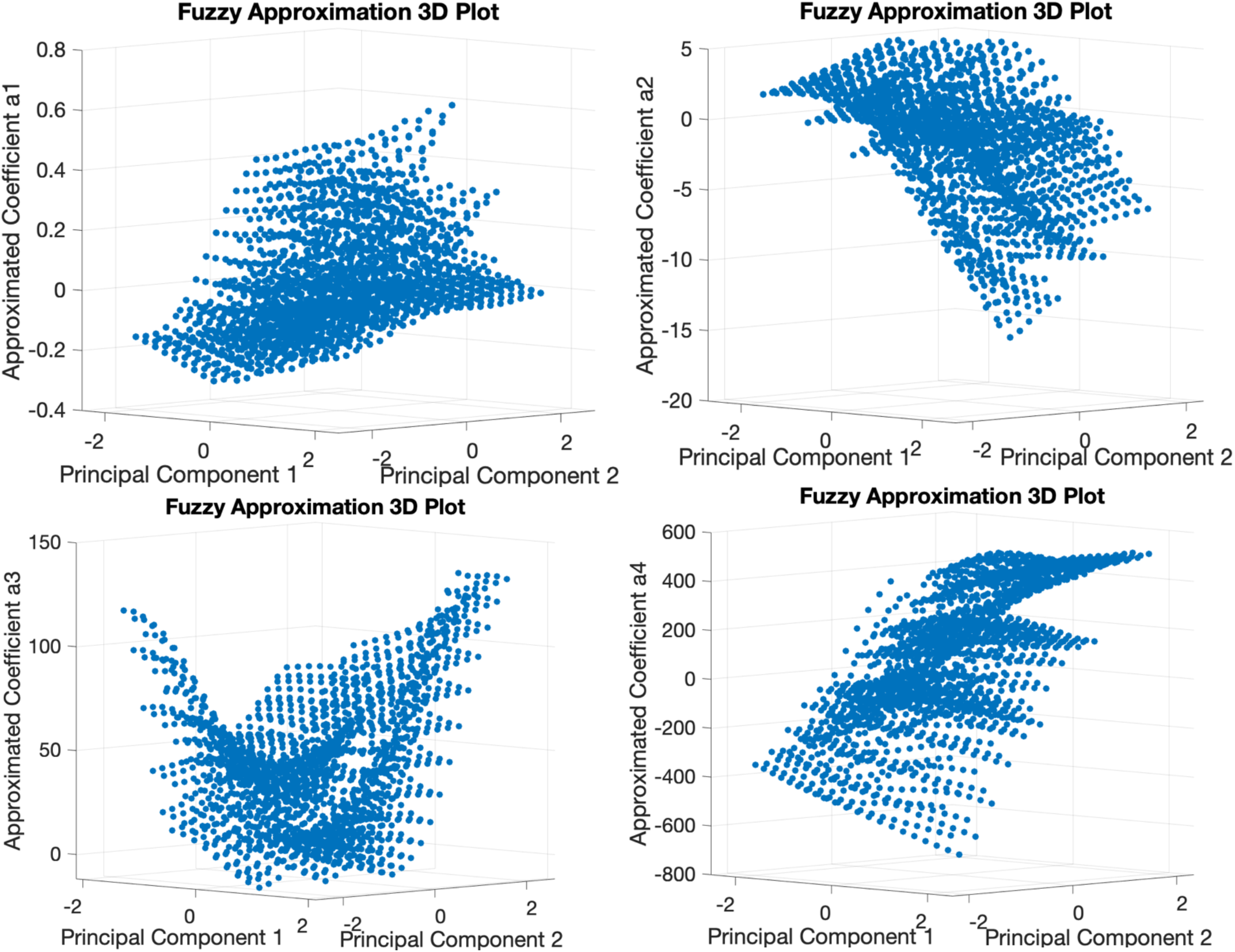
Approximated Fuzzy system output. The figure shows the 3D output of the ANFIS for polynomial coefficients (*a*_1_, *a*_2_, *a*_3_, *a*_4_) and PC1/PC2.

**Supplemental Figure 3.**
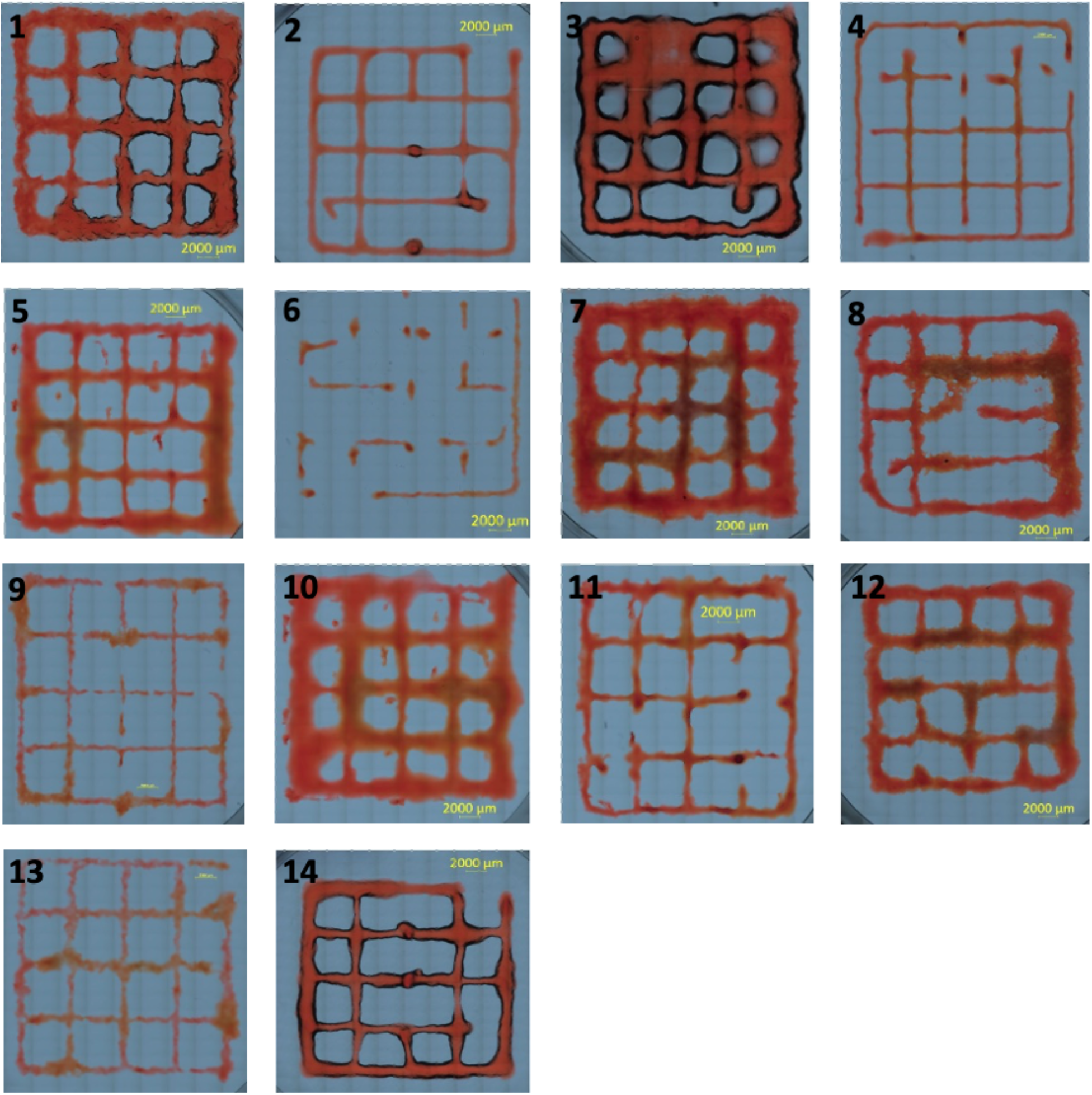
Experimented training dataset. The microscopic images 1-14 represent the extruded training dataset according to the Table 2.

